# Relevance of Antibodies against the Chicken Anaemia Virus

**DOI:** 10.1101/2020.12.15.422992

**Authors:** Max Ingberman, Luiz Felipe Caron, Fernanda Rigo, Liliam C. Araujo, Marco A.P. de Almeida, Letícia Dal Bérto, Breno Castello Branco Beirão

**Affiliations:** Imunova Análises Biológicas, Curitiba, 80215-182, Brazil; Universidade Federal do Paraná, Setor de Ciências Biológicas UFPR, Curitiba, 81531-980, Brazil; Hendrix Genetics, Estr. Mun. Slt-161, Km 08, 53, Salto, 13328-400, Brazil; Elanco Brasil, Av. Morumbi, 8264, São Paulo, 04703-002, Brazil

**Keywords:** Chicken anaemia virus, Maternal antibodies, Vaccination, Breeder, Correlates of protection

## Abstract

**Background:** Chicken Infectious Anaemia (CIA) Virus (CAV) curtails the function of multiple immune compartments. Mortality due to blatant infection is controlled in broilers by passive immunization derived from vaccinated breeders. Therefore, chicks are often assessed by serology to determine maternally-derived antibodies (MDA).

**Methods:** A vaccine overdose-induced model of CIA. The model replicated the most common features of the disease. This model was used to determine the role of MDA in the protection of chicks. Hatchlings were tested for anti-CAV by ELISA and were sorted into groups based on antibody levels. SPF chicks were used as a no-antibody control.

**Results:** Lower specific antibody levels seemed to facilitate viral entry into the thymus, but viral levels, CD4^+^ and CD8^+^ counts, thymus architecture, and haematocrit were preserved by MDA, regardless of its levels.

**Conclusion:** Levels of MDA are not correlated with CIA, but are important for CAV infection.

**IMPORTANCE:** Vaccination is paramount in broiler production. Many of the vaccines are given to broiler breeders, instead of to the broilers themselves. This is cost-effective and practical, since in vaccinating one breeder hundreds of broilers are born with maternally-derived protection. To assess the quality of maternal immunity, antibodies are measured in their chicks. For Chicken Anaemia, this does not seem to suffice to verify protection. This viral disease is very common, and measuring maternal immunity against it determines whether to purchase chicks from a breeder farm. In this study, we verified that antibodies are not correlated with protection from the disease, and therefore should not be used as the sole parameter in assessing immunity against Chicken Anaemia in broilers.

## 1. Introduction

Chicken infectious anaemia is prevalent worldwide in commercial production settings. Nevertheless, clinical disease in broilers has been controlled by the immunization of breeders [1]. The disease is caused by the Chicken Anaemia Virus (CAV), of the Gyrovirus genus. Infection of chicks results in viral replication in haemocytoblasts and lymphoid progenitors, leading to erythrocyte and lymphocyte loss [2,3]. After the first 21 days post-hatch, healthy birds become virtually resistant to clinical disease from CAV infection [3]. Therefore, prevention of the disease is focussed mostly on the vaccination of breeders. It is thus expected that antibodies can be produced and transmitted to their progeny via yolk. In this way, antibody levels in chicks are thought to be correlated to protection against CAV [4].

Here, a practical chicken anaemia disease model was established, demonstrating evident clinical and pathological signs. Using this model, it was possible to verify the role of maternally derived antibodies in the protection of the progeny.

## 2. Materials and Methods

### 2.1. Trials and challenge

Two trials were performed. The first trial aimed to standardize a vaccine overdose trial for chicken anaemia. Previous work demonstrated that immunosuppressed chickens had vaccine virus persistency and showed symptoms, but a simple and reproducible model for chicken anaemia had not yet been established [5]. Here, birds were challenged 1-day post-hatch by 100 times overdose of a CAV vaccine applied via the subcutaneous route with a G24 needle (AviPro™ Thymovac, Elanco, Germany, Lot 07/2019). Parenteral infection is known to induce more prominent Chicken Anaemia disease [6]. This trial was comprised of the following treatment groups: 1) Not challenged, maternally derived antibodies (MDA) – chicks from immunized breeders, not challenged; 2) Challenged, no MDA – SPF chicks, challenged; 3) Challenged, MDA – chicks from immunized breeders, challenged. Chickens were vaccinated at 12 weeks of age with a commercially available product (AviPro™ Thymovac, Elanco, Germany, Lot 05/2017). Chicks were derived from 34-week old breeders. Breeder hens were maintained in Domelia, Brazil. Chicks were hatched in Salto, Brazil. SPF chicks were incubated in the laboratory and were housed in isolators when hatched, when the challenge was performed (SPF birds from Valo BioMedia do Brasil, Brazil). Animals were assessed on D1, D7, D14, D21 for anti-CAV antibodies, virus detection in thymus by qPCR, peripheral blood immunophenotype, and haematocrit measurement. On days 7, 14, and 21 thymuses were collected for histology assessment of lesions.

The second trial aimed to determine the role of maternal antibodies in the protection induced by breeder vaccination. Chickens were vaccinated at 13 weeks of age with a commercially available product (AviPro™ Thymovac, Elanco, Germany, Lot 07/2019). Chicks were derived from 36-week old breeders. Breeder hens were maintained in Itapetininga, Brazil. Chicks were hatched in Salto, Brazil. SPF chicks were incubated in the laboratory and were housed in isolators when hatched, and when the challenge was performed (SPF birds from Valo BioMedia do Brasil, Brazil). SPF birds were used for the infection control since all breeders are vaccinated for CAV in Southern Brazil, thus not being possible to use their offspring as the control group in this study. Chicks from vaccinated breeders were selected for high or low antibody levels following an initial screening by ELISA. Therefore, the experimental groups were: 1) Challenged, Low MDA – chicks from immunized breeders, low levels of anti-CAV antibodies, challenged; 2) Challenged, High MDA – chicks from immunized breeders, high levels of anti-CAV antibodies, challenged; 3) Challenged, no MDA – SPF chicks, challenged. Animals were assessed on D1, D3, D14, D21 for anti-CAV antibodies, virus detection in thymus by qPCR, and haematocrit measurement. On day 14 thymuses were collected for histology assessment of lesions.

In both trials, blood was collected from the wing vein. *Gallus gallus* were from Hendrix Genetics. Organs were collected following culling by cervical dislocation by trained personnel.

The use of animals followed international welfare standards and was approved by the Committee for the use of animals in research of Imunova Análises Biológicas (committee n° 01250.026160/2018-37 (586) registered by the Brazilian Ministry of Science and Technology). Chicks were housed in HEPA-filtered isolators with paper bedding. Handling was performed through gloves that entered the isolators and materials entering or leaving the isolators had to pass through a disinfectant solution. Water and food were supplied *ad libitum*. Feed was commercially formulated, providing for the dietary demands for the age and bird strains used [7]. Water was derived from the public supply. 30 birds were housed in each 1.2 m^2^ isolator. Eight animals per group were removed at each sampling point. Sampling on D1 was conducted before housing the animals.

### 2.2. Laboratory methods

Immunophenotyping was performed by flow cytometry. A no-lysis-no-wash protocol was used for flow cytometry analysis of whole blood, as previously described [8]. Anti-chicken CD4 was FITC-coupled (clone CT-4). Anti-CD8α was PE-coupled (clone CT-8). Anti-chicken CD45 was SPRD-coupled Antibodies by Southern Biotechnology. Absolute cell counts were performed with CountBright (ThermoFisher Scientific, Waltham, MA). Samples were triple stained with the antibodies. “Fluorescence-minus-one” controls were used to diminish spectral overlap. Samples were read in a FACScalibur cytometer (Becton Dickinson, Franklin Lakes, NJ).

For histologic lesion evaluation, thymuses were fixed in formalin. Paraffin-embedded sections were stained with haematoxylin-eosin. A trained veterinary pathologist was responsible for scoring the lesions. They were characterized from 0 to 3, including intermediary scores (0,5). The factors used for grading were medullar/cortical distinction, thymocyte depletion, presence of coagulative necrosis, number of macrophages or segmented cells, and hyperaemia or haemorrhage.

qPCR for virus detection was performed as previously described. The number of plaque-forming units (PFU) was estimated by extracting DNA from viral cell cultures. Positivity for CAV was based on the limit of detection of the method (1054 PFU) [9]. Haematocrit was determined by the microhematocrit capillary technique. A commercially available kit was used for CAV serology, following the instructions of the manufacturer (99-08702, Idexx, Westbrook, ME). Sera were tested at 1:10 and 1:100 dilutions.

### 2.3. Statistical analysis

For statistical analysis, each animal was the experimental unit. Data was assessed based on single or multiple measures and their distribution. Repeated measurements were assessed by two-way ANOVA with Tukey’s posthoc test. Non-parametric data were analyzed by Kruskal-Wallis or Mann-Whitney’s test. Significance was set at *P* < 0.10. Graphs and statistical analyses were performed on GraphPad Prism 6 (GraphPad Software, Inc., La Jolla, CA).

## 3. Results

### 3.1. Establishment of a vaccine-induced model of Chicken Anaemia

A 100 × overdose of the CAV vaccine administered parenterally to day-old chicks was able to induce typical Chicken Anaemia disease. The infection is known to directly affect T cell numbers, but not B cells. Here, virus was detected in high quantities in thymus in animals without maternal immunity. MDA delayed thymic infection (Fig.1A and Fig. 1B) and reduced lesions (Fig. 1C). Total leukocyte counts were reduced in challenged animals without maternally derived immunity, as were CD45^+^CD4^−^CD8^+^ and CD45^+^CD4^+^CD8^−^ cells.

**Fig. 1.**
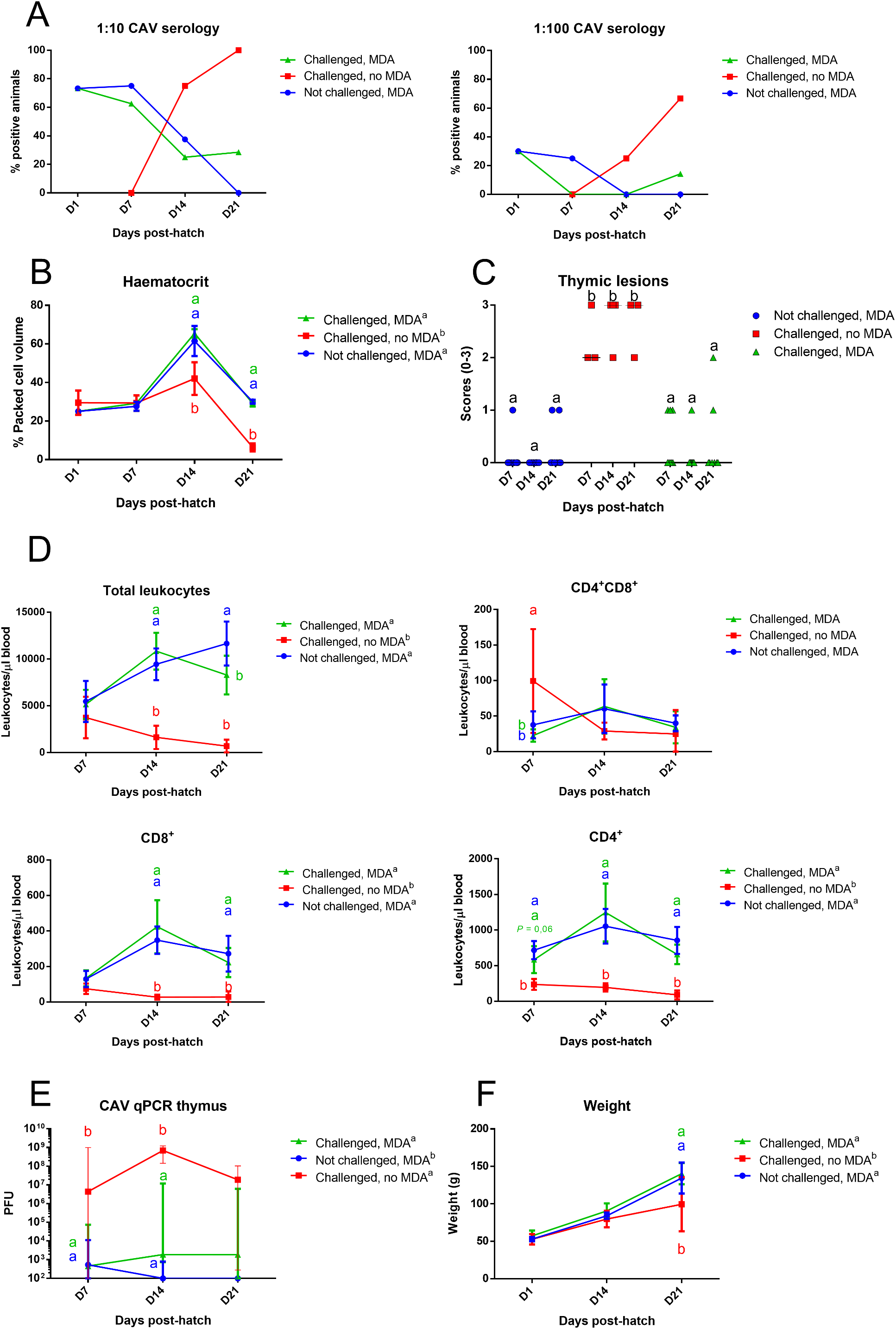
The Chicken Anaemia challenge model was consistent with wild-type infections. Chicks derived from immunized breeders were serologically positive at the start of the trial, whereas SPF birds were induced to produce antibodies by the challenge. Serologic positivity was studied at sample dilutions of 1:10 and 1:100. SPF antibody was not measured in D1 (A). Lack of maternally derived antibodies (MDA) leads to rapid and intense thymus infection following challenge, as measured by qPCR for CAV. MDA partially prevented this (B). Thymic lesions were in accordance with virus levels [n = 4 pools of 2 birds/group] (C). The typical capacity of CAV in affecting progenitor cells seemed to occur, as seen by the reduction in total leukocyte counts and the subsets of CD45^+^CD4^+^CD8- and CD45^+^CD4-CD8^+^, but not in CD45^+^CD4^+^CD8^+^ (D). Similarly, the blood packed cell volume (PCV) was reduced following the challenge (E). Finally, bird body weight was negatively affected by the challenge in the absence of MDA, but not in its presence (F). Statistical analysis by two-way ANOVA with Tukey’s posthoc test (*P* < 0.05 unless stated otherwise), except for histology scores (Kruskal-Wallis test). Significant differences on a given date are indicated by different letters in the colours of the corresponding group. Significant differences throughout the experiment (main treatment effect) are indicated by different letters next to the legend of each graph. The qPCR graph shows the median ± ranges. The histology graph indicates the median, and each dot represents an animal. Other graphs show average ± SD.

Interestingly, triple stained CD45^+^CD4^+^CD8^+^ cells were not affected by the challenge (Fig. 1D). Erythrocyte packed-cell volume was similarly affected by the vaccine overdose (Fig. 1E). Finally, the experimental infection reduced total body weight in chicks without maternal immunity by D21 post-hatch (Fig. 1F).

### 3.2. CAV antibody levels in chicks determine early viral prevalence but not pathogenesis

The established model of Chicken Anaemia was used to determine the role of maternally derived antibodies in the protection conferred to chicks following breeder vaccination. For this, hatchlings were segregated based on their maternal antibody levels (Fig. 2A). Chicks with high MDA had lower prevalence levels of thymic CAV on days 4 and 14 post-hatch (Fig. 2B). Nevertheless, regardless of MDA levels, only on D21 did chicks with MDA reach the same CAV infection levels of SPF birds (Fig. 2B). This was reflected also in lower scores of thymic lesions in chicks that received maternal immunity, regardless of antibody levels (Fig. 2C). Further, erythrocyte packed cell volume was preserved in birds with MDA, regardless of level, whereas SPF birds showed a marked decrease in its levels (Fig. 2D).

**Fig. 2.**
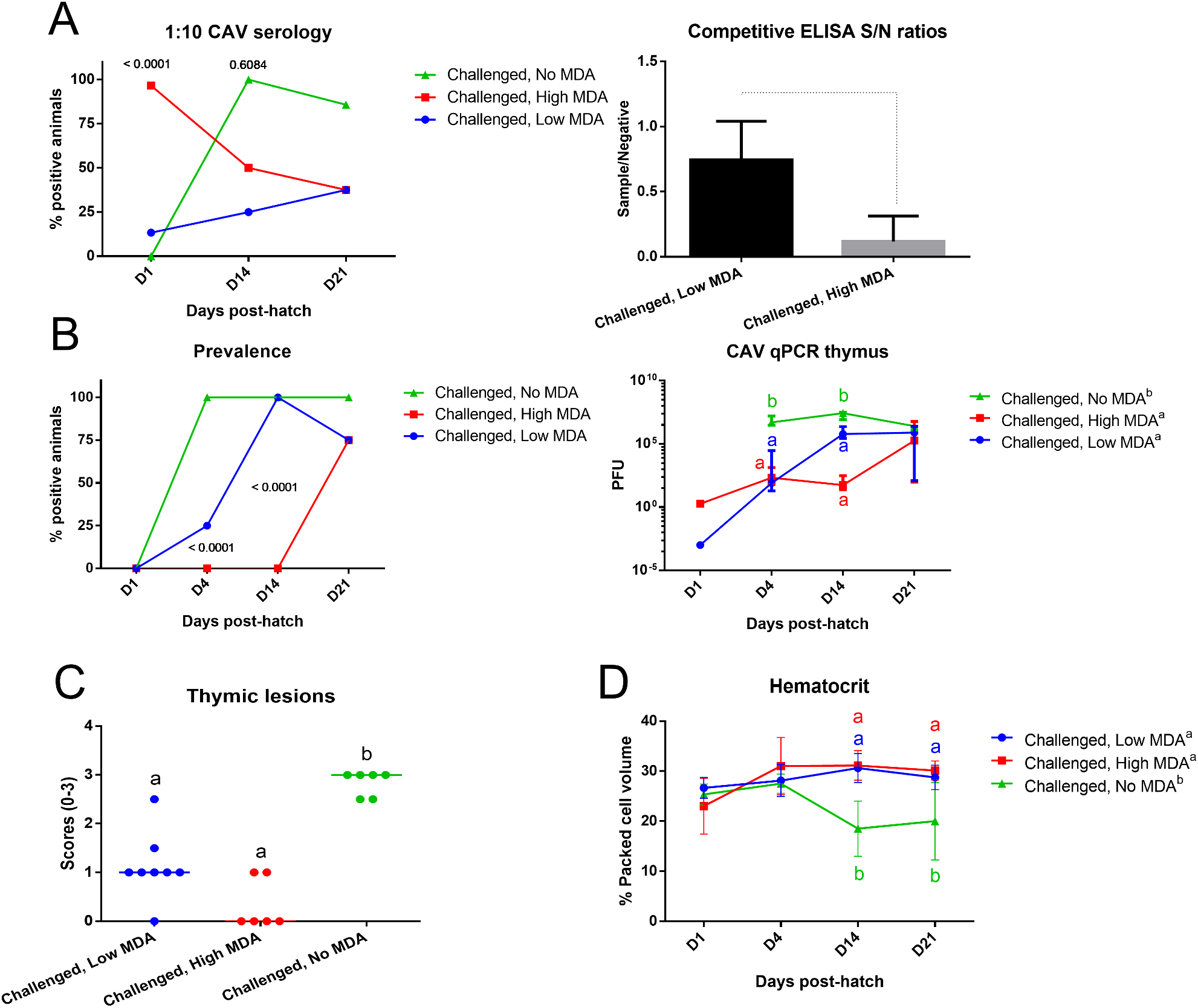
Protection from Chicken Anaemia disease is not dependent on maternally derived antibodies (MDA). Day-old chicks were segregated in isolators based on their antibody positivity and levels for CAV (A). SPF birds were used as MDA-free controls. All animals were challenged with CAV. Higher maternal antibody levels withheld the incidence of CAV in thymuses, but overall viral loads were similar in chicks from vaccinated breeders, regardless of MDA (B). In accordance, thymic lesions (C) and erythrocyte packed cell volume (D) were preserved in the presence of maternal immunity, regardless of the levels of anti-CAV antibodies. Statistical analysis by two-way ANOVA with Tukey’s posthoc test (*P* < 0.05 unless stated otherwise), except for histology scores (Kruskal-Wallis test). Significant differences on a given date are indicated by different letters in the colours of the corresponding group. Significant differences throughout the experiment (main treatment effect) are indicated by different letters next to the legend of each graph. The qPCR graph shows the median ± ranges. The histology graph indicates the median, and each dot represents an animal. Other graphs show average ± SD.

## 4. Discussion

Chicken anaemia virus (CAV) is a common pathogen in commercial farms of birds [1,10]. The virus affects precursor cells in the bone marrow and thymus, impacting on the production of T lymphocytes and erythrocytes, among other cell lineages. Appearance of disease following CAV infection is mostly restricted to the first two weeks post-hatch [3]. Therefore, it has become the norm that immunity is not conferred to chicks by direct vaccination of these birds, but of their breeders [1]. Because of this practice, maternally derived antibody (MDA) levels of day-old chicks are often seen as the ideal correlates of protection [4]. The results of this study suggest that anti-CAV antibody levels in chicks are not associated with disease susceptibility, although there may be a correlation with viral spread.

A challenge model was established for CAV disease, based on the overdosage of a live vaccine. The use of experimental challenge models is relevant since it allows the replication of trials with reproducibility. We also propose the use of a live vaccine overdosage as a CAV challenge model due to the added advantage of enabling laboratories that do not have the expensive virology infrastructure to study the disease. The model was able to replicate the effects of virulent CAV infections in unprotected chicks. During infection, the virus is intensely located in the cortex and corticomedullary junction of the thymus, inducing CD4^+^ and CD8^+^ cell loss and T cell-dependent immune suppression [11,12]. Leukopenia (with loss of CD4^+^ and CD8^+^ cells), anaemia, thymic lesions, and reduced weight gain were induced in challenged, non-immune chicks. Virus could be detected by qPCR in thymus. Passively immunized animals had reduced effects of the challenge, whereas the negative control remained free from alterations. Therefore, the live vaccine experimental model could replicate the effects of wild-type CAV infection and is amenable to replication by laboratories that do not possess virology infrastructure.

The vaccine-induced challenge model was used to determine the effect of maternally derived antibodies in the protection of the progeny. Day-old chicks were grouped based on their anti-CAV levels. As it was expected, SPF birds, that do not carry maternal immunity to CAV, were very susceptible to the challenge, being rapidly infected by the virus and demonstrating anaemia and impairment of immunity. Interestingly, all the chicks derived from vaccinated breeders were consistently protected from the challenge, regardless of the levels of maternal antibodies. Lower levels of MDA increased the early (D4 and D14 post-hatch and challenge) prevalence of CAV, but this did not result in higher average virus levels or in loss of immune cells and erythrocytes. Virus loads in immune organs are correlated to the reduction of tissue lymphocyte counts [6], as observed here with virus quantification in thymus by qPCR, thymic lesions, and numbers of CD4^+^ and CD8^+^ cells.

In conclusion, these findings confirm that maternal immunity is critical in the protection of chicks against CAV disease, but that it is not necessarily mediated by CAV-specific antibodies [6]. Other mechanisms can be involved in the protection of the offspring by maternal immunity: 1) Breeders also transfer high levels of natural antibodies (NAb) to their progeny. These are important components of innate recognition of pathogens in the absence of previous stimulation by an antigen [13]. Immunization alters the NAb repertoire, albeit not specifically against an antigen, as is normally intended by vaccination, due to the unspecific nature of NAbs [14]. In this way, NAbs may partly respond for the improved CAV resistance of chicks with maternally derived immunity; 2) CAV infection induces immune suppression by TCR dysregulation, and maternally derived immune cell effectors may be part of the protection provided by the breeders to their offspring [15]; 3) Maternal influence on the immunity of chicks can also be mediated by hormones passed through the egg [16]. This, and the obvious difference in the genetic background, may have accounted for some of the differences between SPF and commercial birds [17].

## Authors’ contributions

M.I., B.C.B.B., F.R., and L.F.C. were responsible for the trials. M.I., B.C.B.B., L.F.C., L.D.B., and L.C.A. designed the experiment. B.C.B.B. drafted the manuscript and all authors were responsible for its correction and approval.

## Abbreviations

CAV: chicken anaemia virus
D: days
MDA: maternally derived antibodies
PFU: plaque-forming unit
qPCR: quantitative real-time polymerase chain reaction
SPF: specific-pathogen-free

